# Subtractive CRISPR screen identifies factors involved in non-canonical LC3 lipidation

**DOI:** 10.1101/2020.11.18.388306

**Authors:** Rachel Ulferts, Elena Marcassa, Lewis Timimi, Liam C Lee, Andrew Daley, Beatriz Montaner, Suzanne D. Turner, Oliver Florey, J. Kenneth Baillie, Rupert Beale

**Affiliations:** The Francis Crick Institute, London, UK; Department of Pathology, Cambridge University, Cambridge, UK; Signalling Programme, Babraham Institute, Cambridge, UK; Edinburgh University, Edinburgh, UK

**Keywords:** non-canonical autophagy, CRISPR, ATG16L1, RalGAP, vATPase

## Abstract

Although commonly associated with autophagosomes, LC3 can also be recruited to membranes in a variety of non-canonical contexts. These include responses to ionophores such as the M2 proton channel of influenza A virus. LC3 is attached to membranes by covalent lipidation that depends on recruitment of the ATG5-12-16L1 complex. Non-canonical LC3 lipidation requires the C-terminal WD40 domain of ATG16L1 that is dispensable for canonical autophagy. We devised a subtractive CRISPR knock-out screening strategy to investigate the requirements for non-canonical LC3-lipidation. This correctly identified the enzyme complexes directly responsible for LC3-lipidation. We additionally identified the RALGAP complex as important for M2-induced, but not ionophore drug induced LC3 lipidation. In contrast, we identified ATG4D as responsible for LC3 recycling in M2-induced and basal LC3-lipidation. Identification of a vacuolar ATPase subunit in the screen suggested a common mechanism for non-canonical LC3 recruitment. Influenza-induced and ionophore drug induced LC3-lipidation leads to association of the vacuolar ATPase and ATG16L1 and can be antagonised by Salmonella SopF. LC3 recruitment to erroneously neutral compartments may therefore represent a response to damage caused by diverse invasive pathogens.

## Introduction

Autophagy is a catabolic process characterised by the delivery of cytoplasmic material to the lysosome for degradation (Mizushima and Komatsu, 2011). A distinct feature of this pathway is the relocalisation of microtubule-associated proteins 1A/1B light chain 3 (LC3). Upon induction of autophagy, LC3 becomes covalently conjugated to phosphatidylethanolamine (PE) at sites forming double-membrane autophagosomes (Mizushima et al., 1998). Lipidation depends on the activity of two ubiquitin-like conjugation systems comprising ATG3, ATG5, ATG7, ATG10 and ATG12 (Kaufmann et al., 2014). ATG5 and ATG12 form a complex with ATG16L1 that catalyses the transfer of activated LC3 to PE, in a manner analogous to the role of an E3 ligase. The localisation of this complex determines site-specificity of LC3 lipidation (Fujioka et al., 2014). The soluble form of LC3 is referred to as LC3-I, and the PE-conjugated form of LC3 as LC3-II.

LC3-II also decorates various single membrane compartments in response to different stimuli (Florey et al., 2011, 2015; Jacquin et al., 2017; Sanjuan et al., 2007). Examples of this non-canonical pathway include LC3-associated phagocytosis (LAP), macropinocytosis and entosis (Florey et al., 2011, 2015). In all of these examples a subset of these endocytic vesicles acquires LC3-II. This ‘non-canonical autophagy’ has also been implicated in the regulation of host homeostasis following infection with Influenza A virus (IAV) (Fletcher et al., 2018). IAV is a negative sense, segmented RNA virus responsible for seasonal flu epidemics and global outbreaks. Upon infection of the cell, the viral M2 protein, a small proton selective ion channel or “viroporin”, promotes LC3-lipidation (Beale et al., 2014; Gannagé et al., 2009; Ren et al., 2016; Zhirnov and Klenk, 2013). LC3-II accumulates at intracellular vesicles and the plasma membrane (Beale et al., 2014). Influenza M2 dissipates intracellular proton gradients, resulting in erroneously neutral compartments (Ciampor et al., 1992; Henkel et al., 1999). LC3-lipidation is dependent on the ion channel activity of M2 (Fletcher et al., 2018; Ren et al., 2016). Additionally, the C-terminal region of the IAV-M2 interacts directly with LC3 through a highly conserved LC3-interacting region (LIR) (Beale et al., 2014). We have recently shown that the molecular mechanism of LC3-lipidation induced by M2 differs from that of canonical, starvation induced LC3 lipidation. In contrast to canonical autophagy, the recruitment of the E3-like ATG12-ATG5/ ATG16L1 complex during M2-induced LC3-lipidation depends on the WD repeat-containing C-terminal domain (WD40 CTD) of ATG16L1 but is independent of FIP200 and WIPI2b binding (Fletcher et al., 2018). The WD40 CTD-dependency of ATG16L1 recruitment is also observed during other non-canonical LC3-lipidation events, including responses to ionophores, LC3-associated phagocytosis (LAP) and entosis (Fletcher et al., 2018). The molecular mechanism of ATG16L1 recruitment to these membranes is unknown, but provocatively WD40 CTD has been reported to interact with the v-ATPase in the context of Salmonella infection. This process is antagonised by the salmonella effector SopF (Xu et al., 2019).

To identify the genes involved in this novel cellular pathway, we performed a genome wide genetic knockout screen using CRISPR Cas9 technology. We exploited the resistance of GFP-tagged LC3-II to removal from permeabilised cells by detergents such as saponin to enable us to sort cells based on fluorescence (Eng et al., 2010). To discriminate between true hits and genes that affect expression of the GFP-LC3 marker, we also performed the screen without permeabilisation, and subtracted these results on a guide-by-guide basis. This differential screen uncovered several host factors involved in the regulation of M2-dependent LC3 lipidation, including all six components of the core lipidation machinery. We also identified a subunit of the v-ATPase (V0A1) and the GTPase activator RALGAP beta subunit as required for optimal LC3 lipidation. Conversely, deletion of the cysteine protease ATG4D enhanced LC3 lipidation and accumulation in cells expressing IAV M2.

## Results

### The proton channel activity of M2 is required for M2-induced LC3-lipidation

We previously showed that Influenza A virus (IAV) induces LC3 lipidation and relocalisation to the plasma membrane through a pathway that differs from canonical autophagy (Beale et al., 2014). This event is dependent on the proton channel activity of the M2 protein and WD repeat-containing C-terminal domain (WD40 CTD) of ATG16L1 (Fletcher et al., 2018).

Other WD40 CTD -dependent LC3-lipidation processes target endolysosomal vesicles and thus components overlap with the pathway required for IAV entry into the cell (Florey et al., 2011; Sanjuan et al., 2007). We therefore reasoned that this potential overlap between M2-induced LC3-lipidation and virus entry might obscure target hits in a screen using virus infection. Alternatively, expression of IAV M2 has been shown to be sufficient for the induction of proton channel-dependent LC3-lipidation in the absence of other viral components (Ren et al., 2016; Zhirnov and Klenk, 2013). We therefore established a doxycycline-inducible M2-expression cell line. We chose the M2 protein of IAV strain A/Udorn/72 as its proton channel is sensitive to inhibition by amantadine. Expression of M2 led to LC3-relocalisation and lipidation similar to that observed after infection with IAV (Figure 1A and B). Relocalisation was inhibited by amantadine in M2-expressing and MUd (a reassortant strain of IAV A/Rico/8/34 (PR8) with the M2-encoding segment of strain Udorn) -infected cells but not in cells infected with the amantadine resistant strain PR8, confirming that the proton channel activity of M2 is required for M2-induced LC3-lipidation (Figure 1A and C).

**Figure 1.**
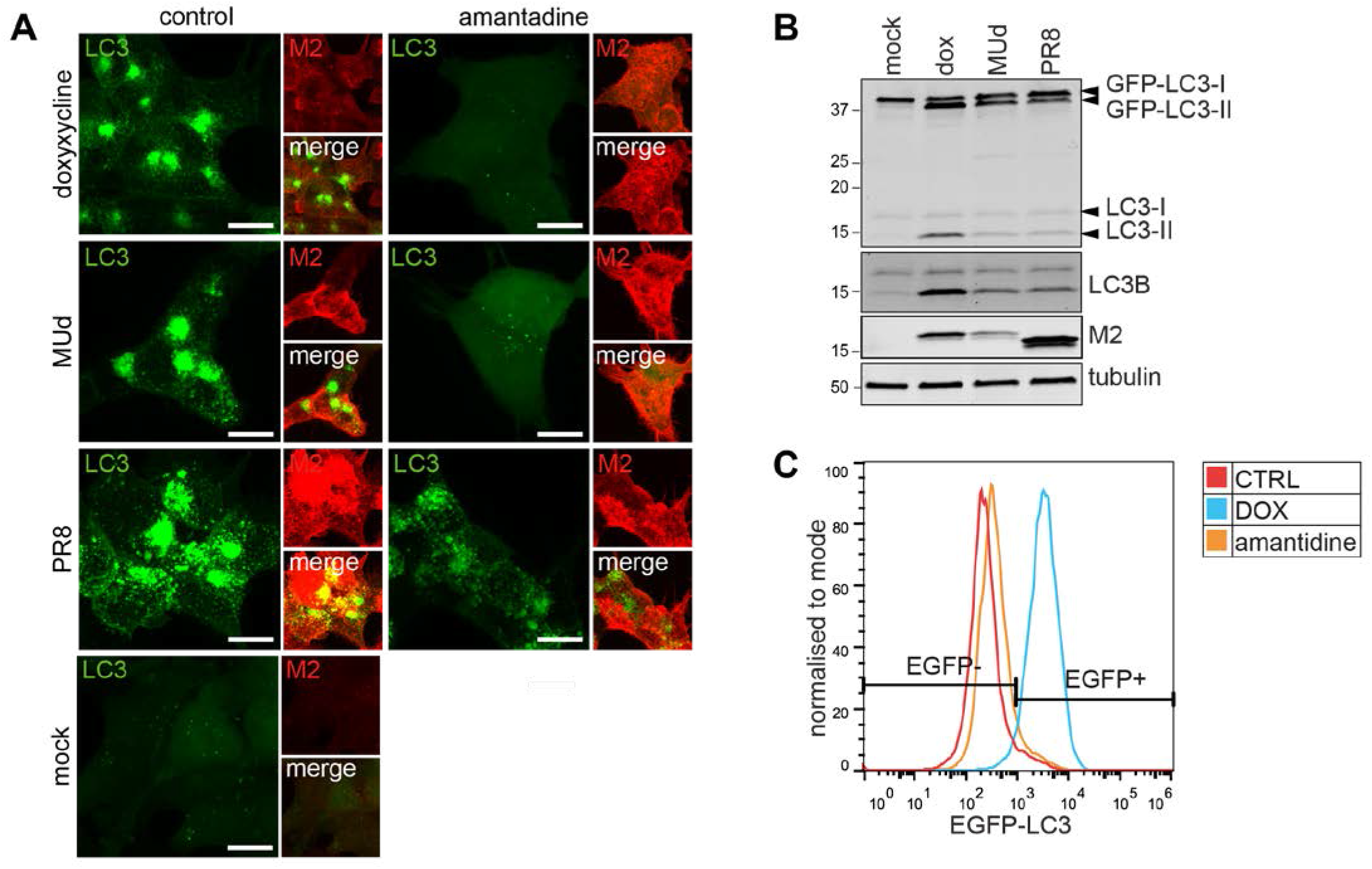
M2 proton channel activity is sufficient for LC3B lipidation. A) HCT116 EFGP-LC3B TetON M2 cells treated for 16 h with 3 μg/ml doxycycline or infected with IAV PR8 or MUd at an MOI of 3 PFU per cell, with or without 5 μM amantadine. Cells were then fixed and stained for M2 protein. Scale bar 10 μM. B) Western blot analysis of LC3 lipidation in HCT116 EFGP-LC3B TetON M2 cells treated for 16 h with 10 μg/ml DOX or infected with IAV PR8 or MUd at an MOI of 3 PFU per cell. C) FACS analysis of levels of membrane associated EGFP-LC3B. HCT116 EFGP-LC3B TetON M2 cells were treated for 16 h with 10 μg/ml DOX, with or without 5 μM amantadine.

### A genome wide CRISPR-Cas9 screen identified factors involved in M2-induced LC3-lipidation

To identify factors involved in this pathway we performed a CRISPR-Cas9 screen in HCT116 EGFP-LC3 cells expressing the IAV-M2 protein under a doxycycline inducible promoter. Briefly, cells were transduced with the GECKO library V2 (Sanjana et al., 2014) encoding 6 gRNAs per gene. Cells were expanded for 10 days post-transduction to maximise gene editing. M2 expression was induced for 16 hours with doxycycline. Permeabilisation of cells with saponin was used to wash away unlipidated LC3 as described in Eng et al., 2010. Since EGFP fluorescence is the read-out, genes involved in EGFP expression from the MMLV based promoter as well as genes involved in LC3-lipidation in this context would be expected to be identified using this method. The pool of cells was therefore split into two, either permeabilised with saponin or not, stained for M2 expression and sorted by FACS according to the top and bottom 10% EGFP fluorescence levels. (Figure 2A). After purification of genomic DNA and sequencing, analysis was performed as per Li et al., 2020. Genes directly responsible for LC3-lipidation were identified in the saponin permeabilised set. Genes such as FAM208A, a component of the HUSH complex, and MAX (MYC Associated Factor X), a known transcription factor, were identified in both the permeabilised and unpermeabilised screens (figure 2B and supplementary figure 1). This is expected since the MMLV promotor, which drives GFP-LC3 marker expression in HCT116 EGFP-LC3 cells, can be silenced by the HUSH complex (Tchasovnikarova et al., 2015). In a z-z scatter plot they therefor align along the z=y axis (Figure 2B). To isolate the genes responsible for promoting or antagonising non-canonical autophagy, we subtracted the Z scores of each guide in the non-permeabilised dataset from the corresponding guides in the permeabilised dataset (supplementary figure 1). This showed the top 6 hits as required for LC3-lipidation to be ATG7, ATG12, ATG16L1, ATG5, ATG10 and ATG3 (Figure 2C and supplementary figure 1A). This subtractive screening approach therefore correctly identified all components of the core LC3-lipidation machinery. Two other interesting candidate genes were ATP6V0A1 and RALGAPB (figure 2D), both of which were analysed further. Only one gene, ATG4D, was strongly identified as a potential antagonist of LC3-lipidation (figure 2D).

**Figure 2:**
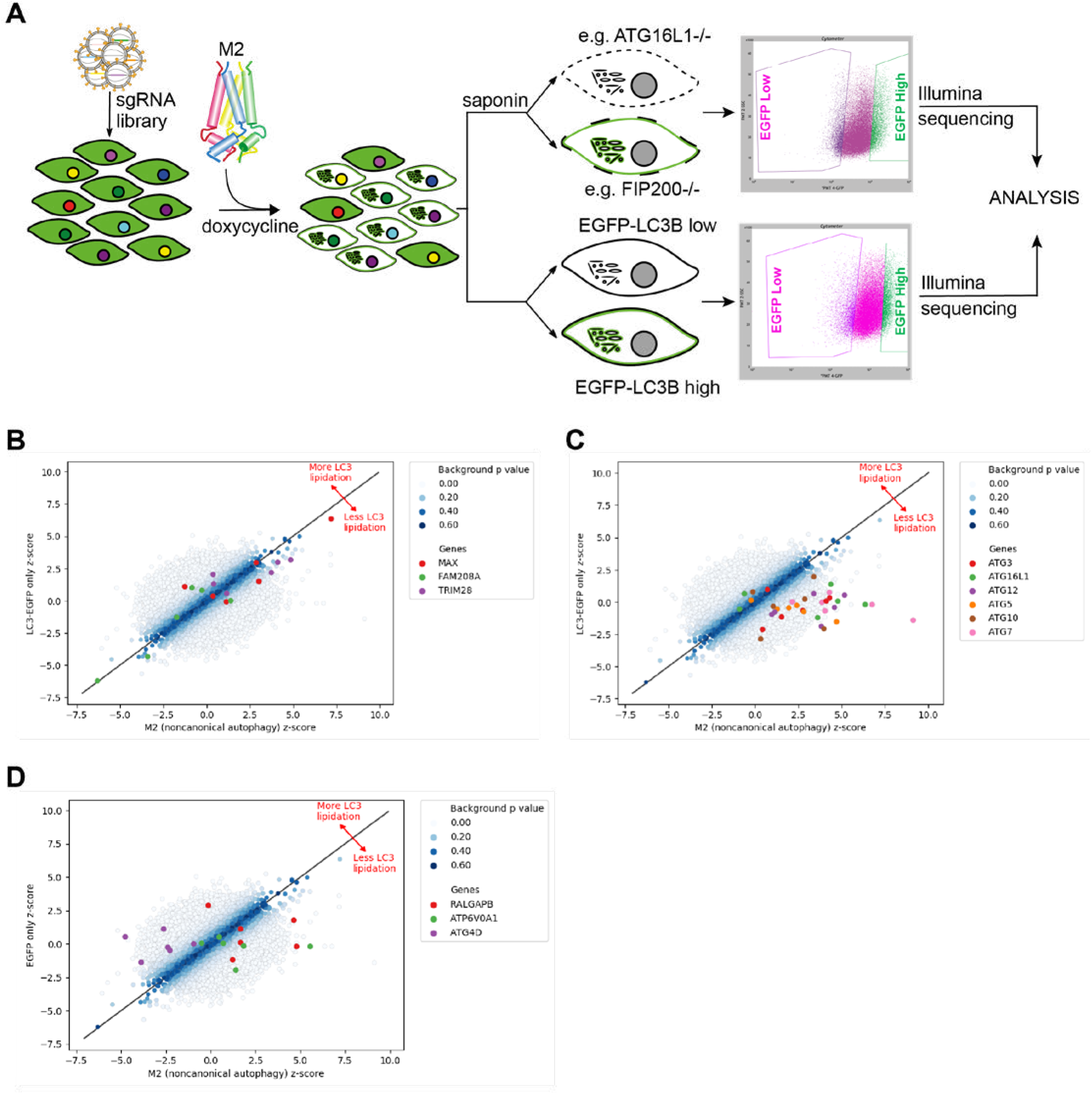
Substractive CRISPR screen identifies factors involved in non-canonical LC3 lipidation. A) Schematic depiction of the CRISPR screen. M2 expression was induced in library transduced cells and the top and bottom 10% EGFP expressing cells sorted in permeabilised M2 expressing cells (top) and in unpermeabilised cells. SgRNA representation was determined by Illumina sequencing and factors involved in M2-induced LC3 lipidation and were identified by substractive comparison of permeabilised and unpermeabilised treatment conditions. B, C, D) Scatter plots showing the representation of sgRNAs B) affecting EGFP-LC3B expression, C) targeting the core LC3-lipidation machinery and D) targeting selected genes required for or counteracting LC3 lipidation.

### The role of the v-ATPase in M2-induced LC3-lipidation

The v-ATPase has been previously implicated in different types of non-canonical autophagy, including entosis, LAP and ionophore-induced LC3-lipidation, on the basis of their sensitivity to the v-ATPase inhibitor bafilomycin A1 (Fletcher et al., 2018; Florey et al., 2015). Recent findings have revealed a requirement for the interaction of the v-ATPase complex with ATG16L1 in the selective degradation of bacteria (xenophagy) (Xu et al., 2019). In the same study, the authors demonstrated the ability of the Salmonella effector SopF, to interfere with this interaction. In light of this new evidence, we sought to investigate a role for this v-ATPase-ATG16L1 axis in the regulation of non-canonical autophagy, including M2-induced lipidation. In common with Xu et al., we failed to obtain knock out cell lines, consistent with v-ATPase function being required for cell viability. Following infection with IAV strain PR8, the V1A subunit of the v-ATPase co-immunoprecipitated with endogenous ATG16L1 (Figure 3A). The ionophore drug monensin increases levels of LC3-II by both canonical autophagy and WD40 CTD-dependent LC3-lipidation (Fletcher et al., 2018). By inhibiting canonical autophagy with a VPS34 inhibitor, IN-1, we were able to demonstrate the association of ATG16L1 and the v-ATPase in another WD40 CTD-dependent LC3-lipidation context (Figure 3B, reciprocal IP in supplementary figure 2A). This interaction was further confirmed using a panel of different cell lines after treatment with IN-1/monensin (supplementary figure 3B). Similar observations were made in HCT116 ATG16L1 KO cells reconstituted with Flag-S-mATG16L1 following doxycycline-induced M2 expression (supplementary figure 3C). Next, we investigated the effect of expression of the bacterial effector SopF on WD40 CTD-domain dependent LC3-lipidation. As mentioned above, this bacterial effector has been shown to inhibit WD40 CTD-dependent LC3-lipidation of the salmonella containing phagophore (Xu et al., 2019). We generated HCT116-EGFP-LC3B tet-ON M2 cell lines stably expressing mCherry-SopF or mCherry alone as a control. In this system, we were able to compare relocalisation after M2 expression or canonical autophagy induction using the mTOR inhibitor, Torin1. In cells treated with doxycycline, SopF expression inhibited LC3 puncta formation and relocalisation to the plasma membrane (Figure 3C). SopF had no effect on Torin 1 induced LC3 puncta formation (Figure 3C), indicating that the inhibition is specific for non-canonical LC3-lipidation. We confirmed this result by western blot analysis (Figure 3D) and in the context of IAV infection (Figure 3E). Collectively, these data confirm the role of the v-ATPase-ATG16L1 axis in the activation of WD40 CTD-dependent LC3-lipidation.

**Figure 3:**
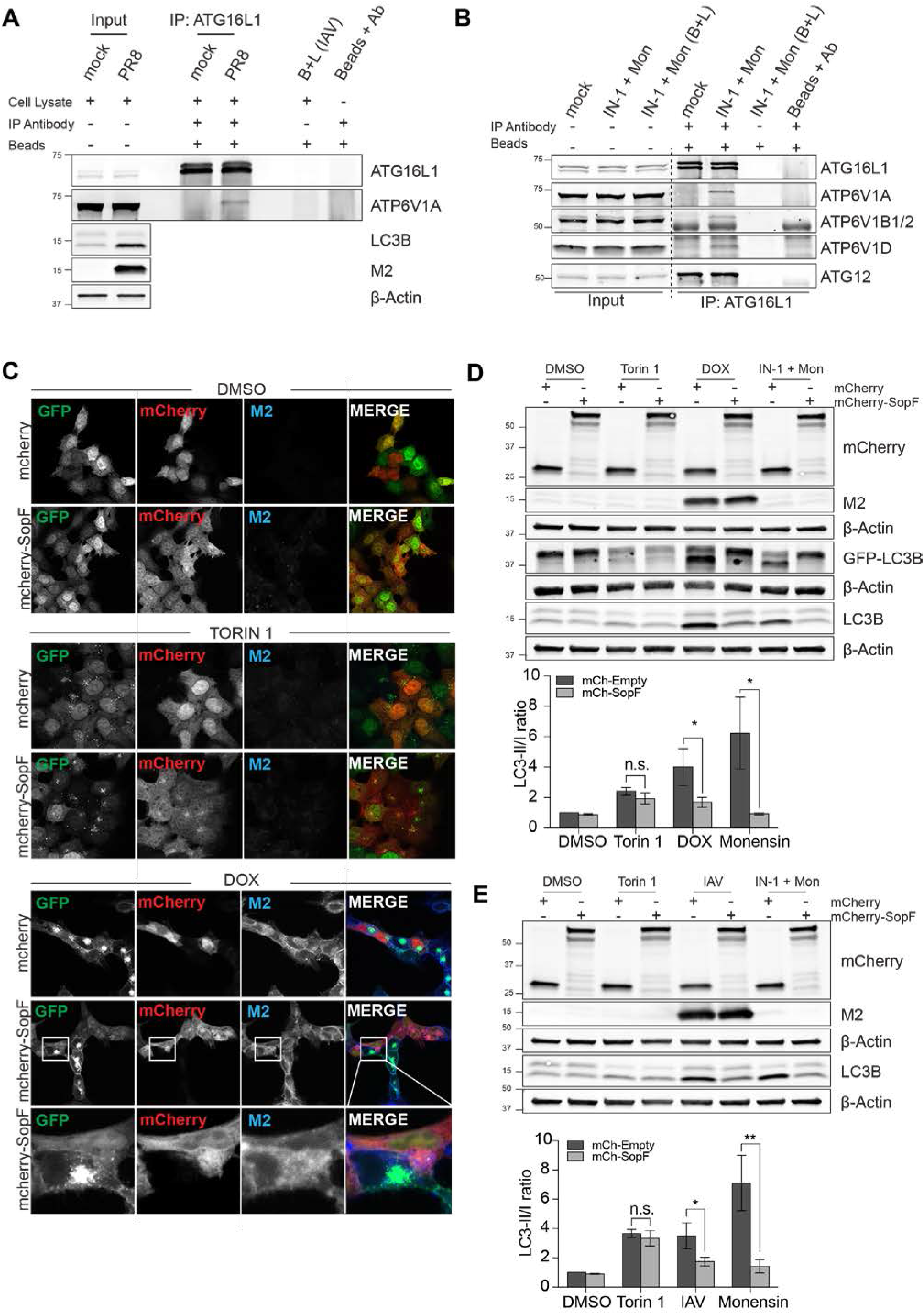
The v-ATPase is required for M2-induced LC3-lipidation. A) Immunoprecipitation analysis of endogenous ATG16L1 in HCT116 cells infected for 16 hrs with PR8. B) Immunoprecipitation analysis of endogenous ATG16L1 in HeLa cells treated with VPS34 IN-1 (pretreatment: 1 μM for 30 minutes) followed by monensin (100 μM for 1 h). C) HCT116 EGFP-LC3B TetON-M2 cells stably expressing mCherry or mCherry-SopF following treatment with Torin 1 (250 nM for 3 h) or doxycycline (10 μg/ml for 16 h) or mock treated. D) LC3B lipidation analysis in HCT116 EGFP-LC3B TetON-M2 cells stably expressing mCherry or mCherry-SopF after treatment with either Torin 1 (250 nM for 3 h), doxycycline (10 μg/ml for 16 h) or VPS34 IN-1 pretreatment (1 μM for 30 minutes) followed by monensin (100 μM for 1 h). The graph shows fold change in LC3II/LC3I ratio relative to mCherry DMSO from three independent experiments. Bars show mean±SD. Unpaired Student’s t-tests were performed. *, P < 0.05. E) HCT116 stably expressing mCherry or mCherry-SopF treated as in B or infected with PR8 for 16 h (MOI 10 PFU per cell). The graph shows fold change in LC3II/LC3I ratio relative to mCherry DMSO from three independent experiments. Bars show mean±SD. Unpaired Student’s t-tests were performed. *, P < 0.05; **, P<0,01.

### The RalGAP complex is important for M2-induced LC3-lipidation

RalGAPβ is the non-catalytic subunit of the Ral GTPase Activating Protein complex and forms a heterodimer with a RalGAPα subunit (Shirakawa et al., 2009). This complex acts as a GTPase activator for Ras-like small GTPases RALA and RALB. To validate RalGAPβ involvement in the regulation of LC3-lipidation we generated HCT116 tet-ON M2 cells RalGAP KO cells using CRISPR-Cas9 and confirmed the absence of RalGAPβ expression by western blot (figure 4A). Depletion of RalGAPβ strongly reduced M2-induced LC3-lipidation (figure 4B) and relocalisation of EG-FP-LC3B (figure 4F).

**Figure 4.**
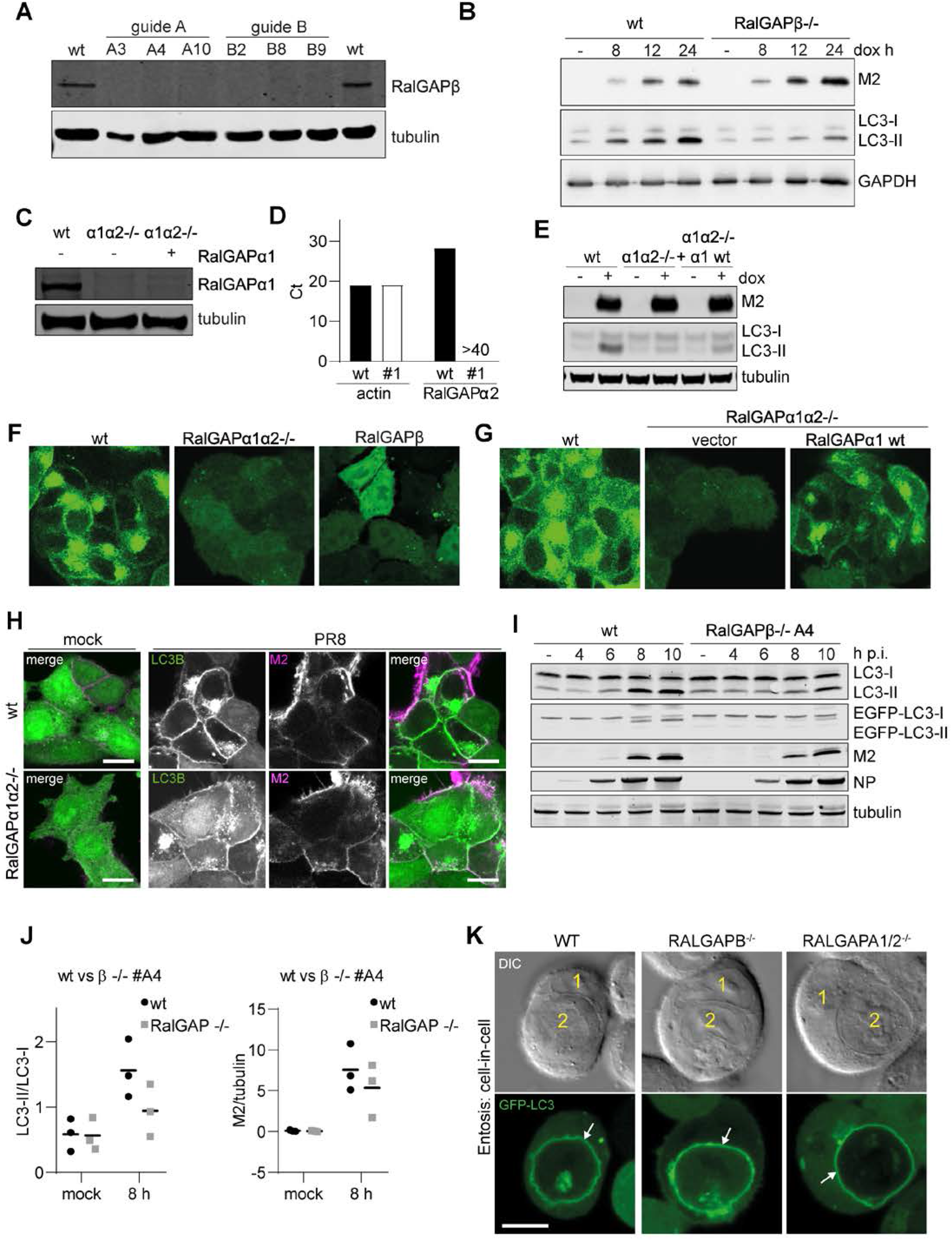
RalGAP depletion inhibits EGFP-LC3B relocation in response to M2 expression but not other forms of non-canonical LC3-lipidation. A) Western blot validation of HCT116 EFGP-LC3B TetON M2 cells knock out (ko) for RalGAPβ. B) LC3 lipidation analysis of HCT116 EFGP-LC3B TetON M2 WT and RalGAPβ−/− ko cells after at the indicated timepoints after induction with 10 μg/ml dox. C) Western blot validation of HCT116 EFGP-LC3B TetON M2 cells ko for RalGAPαlα2. D) qPCR quantification of RalGAPα2 expression in HCT116 EFGP-LC3B TetON M2 cells wt and ko for RalGAPα1α2. E) LC3 lipidation analysis of HCT116 EFGP-LC3B TetON M2 cells WT and ko for RalGAPα1α2 after 16 hr treatment with 10 μg/ml dox. F) HCT116 EGFP-LC3B TetON-M2 cells depleted for RalGAPα1α2 and RalGAPβ after 16 hr treatment with 10 μg/ml dox. G) HCT116 EGFP-LC3B TetON-M2 cells depleted for RalGAPα1α2 were transduced with RalGAPα1 expressing lentivirus or empty vector and M2 expression induced for 16 h. H) HCT116 EGFP-LC3B TetON-M2 cells depleted for RalGAPα1α2 and RalGAPβ and infected with PR8 for 8 hrs. Cells were then fixed and stained for M2 protein. Scale bar 10 μM. I) LC3 lipidation analysis of HCT116 EGFP-LC3B TetON-M2 cells depleted for RalGAPβ (clone A4) after infection with PR8 at on moi of 10 PFU per cell. J) Quantification of H. Graphs show mean of LC3II/LC3I ratio (left) and M2 expression normalised to tubulin (right) from three independent experiments. K) EGFP-LC3B recruitment to entotic vesicles in HCT116 EFGP-LC3B TetON M2 wt, RalGAPα1α2−/− and RalGAPβ−/− cells analysed by fluorescence (top panel) and bright field (bottom panel) live cell microscopy. Scale bar 10 μM.

We also analysed LC3 behaviour in cells depleted of the other subunit of the complex, RalGAPα. Mammalian cells encode two paralogues, RalGAPα1 and RalGAPα2. In human cells, these possess 53% identity across the whole sequence and 83% sequence identity in the GAP domain (Martin et al., 2014). We generated cell clones devoid of expression of both paralogues. Absence of RalGAPα1 expression was confirmed by western blot (figure 4C), while absence of RalGAPα2 expression, due to the lack of a suitable antiserum, was confirmed by qPCR (figure 4D). Lack of the RalGAPα subunits also strongly reduced M2-induced EGFP-LC3B relocalisation (figure 4F) and LC3-lipidation (figure 4E), implicating the RalGAP complex in M2-induced LC3-lipidation.

We then tested the effect of RalGAP knock out in LC3-lipidation during IAV infection. RalGAPβ−/− cells clone A4 and wt cells were infected with IAV strain PR8 and LC3-lipidation at different time points post infection (p.i.) analysed by western blot. While we observed a reduction in LC3-lipidation in RalGAPβ−/− cells, this effect was reduced compared to the M2-expression system (figure 4H and I). We also observed a slight reduction in M2 expression levels (figure 4J). Identical effects on LC3-lipidation concurrent with a slight reduction in M2 expression levels were observed in infection of RalGAPβ−/− clone B8 - which was generated using a different sgRNA to clone A4 - or the RalGAPα1α2−/− cell line (supplementary figure 3). No effect on virus replication was found (supplementary figure 3C). The observed effect was thus specific for the depletion of the RalGAP complex, and not a clonal artefact.

A similar reduction in LC3-lipidation and M2 expression was observed using the chimeric influenza virus MUd (supplementary figure 3B). This indicates that the effect on LC3-lipidation and M2 expression are not due to strain specific differences in the M2 protein sequence.

We have previously shown that the dependence of M2-induced LC3-lipidation on the WD40 CTD of ATG16L1 is shared by other “non-canonical” pathways, namely LAP, entosis and drug-induced endosomal perturbation (Fletcher et al., 2018). Entosis is a process observed in cancer cells by which viable cells become engulfed by another cell. Upon death of the internalised cells the surrounding vesicle membrane of the engulfing cell becomes decorated with LC3 (Florey et al., 2011). To test if RalGAP is important in LC3-lipidation in this process we analysed entotic events in RalGAP−/− and wt cells. No significant difference could be observed, indicating that RalGAP is dispensable for this activity (figure 4K). We also could not observe an effect of RalGAP knock out on LC3-lipidation and EGFP-LC3B punctae formation during monensin-induced endosomal perturbation (supplementary figure 4A and B). In summary, RalGAP does not appear to play a role in other forms of WD40 CTD-domain-dependent LC3-lipidation but to be specific for IAV M2.

To further exclude that the observed effects were due to off target effects of the chosen sgRNAs we expressed RalGAPα1 in RalGAPα1α2−/− cells using lentiviral transduction. Expression levels of RalGAPα1 remained lower than levels in wt cells (figure 4C). This was the case when expressing untagged as well as N-terminally mCherry-tagged RalGAPα1 (data not shown). Use of a CMV promotor driven expression system in place of the PGK promotor did not improve expression levels (data not shown). However, a partial recovery of M2-induced LC3-lipidation and relocalisation was observed (figure 4E and G). In summary, we concluded that the RalGAP complex is important for M2-induced LC3-lipidation.

### ATG4D depletion enhances LC3B lipidation levels

ATG4D was the only gene targeted that appeared to give rise to higher LC3-lipidation in the screen. The ATG4 family comprises 4 isoforms, ATG4A, B, C and D. ATG4C and ATG4D are its least characterised members. During autophagy, ATG4 is required both for the processing of the newly synthesised pro-LC3/GABARAP and for the recycling of the PE-conjugated form to the autophagosomal membranes (Yu et al., 2012). We have shown that LC3 can also become conjugated to phosphatidylserine (PS)-enriched membranes during non-canonical autophagy. ATG4D, but not the other paralogues, appears to be uniquely capable for the de-conjugation of LC3-PS (Durgan et al., 2020).

Depletion of ATG4D in HCT116 cells using siRNA results in a slight increase in LC3-lipidation in fed conditions (figure 5A). Levels of LC3B-II were also higher in IAV-infected and amino-acid starved cells with reduced ATG4D levels compared to LC3B-II levels in control cells. We noticed a similar increase in LC3B-II levels in several clones of HCT116 ATG4D−/− cells (figure 5B and supplementary figure 5A). Levels of M2-induced LC3B-II were significantly higher in ATG4D−/− cells compared to wt (figure 5C).

**Figure 5:**
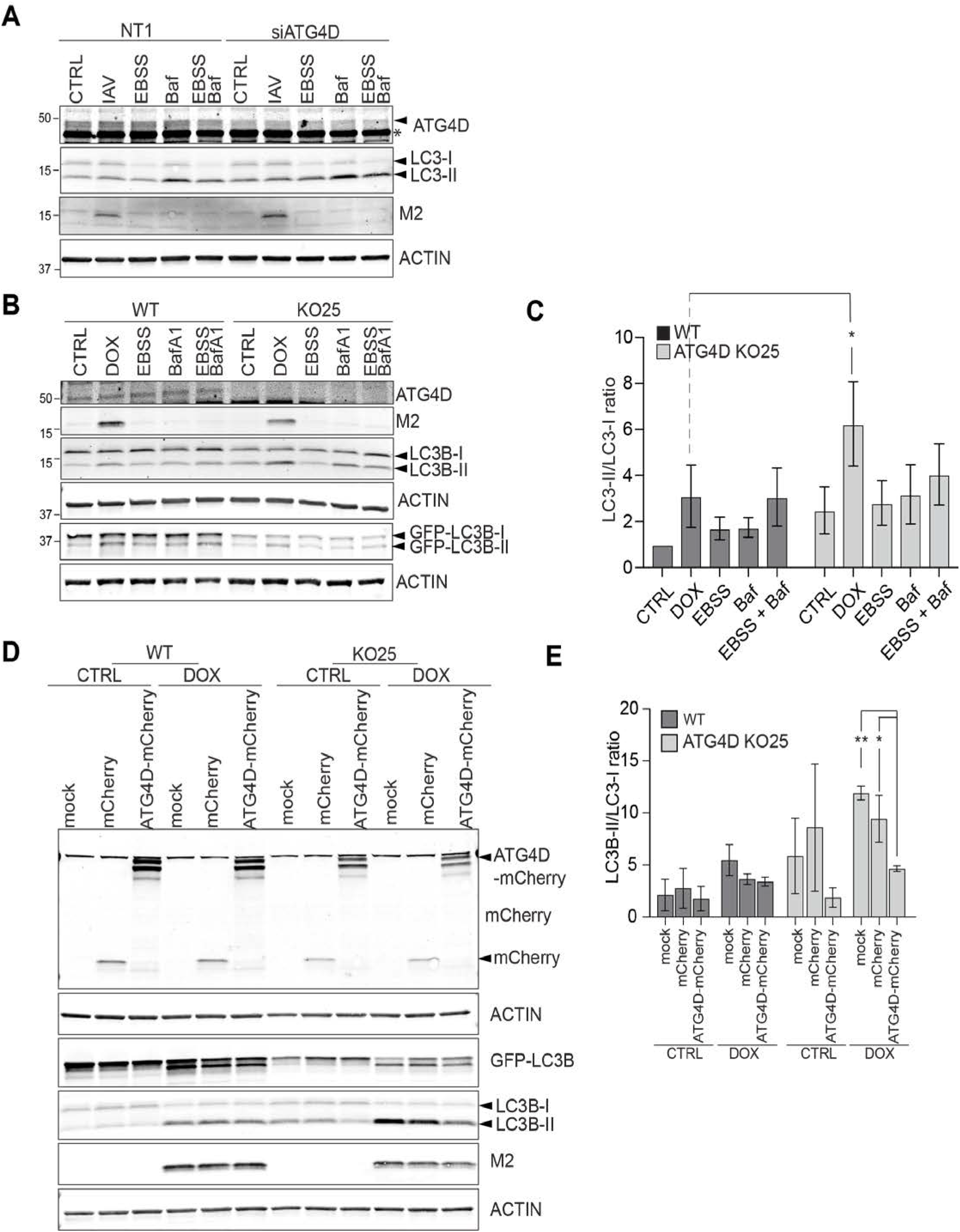
ATG4D depletion enhances LC3B lipidation. A) HCT116 cells were treated with siRNA as indicated and LC3B lipidation was analysed after incubation for 8 hr with PR8 (20 moi), 2 hr with EBSS, 1 hr with 200 nM Bafilomycin A1 and a combination of EBSS together with Bafilomycin A1. arrowhead indicates ATG4D specific band, * background band. B) HCT116 EGFP-LC3 TetOn-M2 WT and ATG4D KO clone 25 were treated with 10 μg/ml dox for 8 h or as in A. C) Quantification of the experiment in B. The graph shows the change in LC3II/LC3I ratio from 4 independent experiments; bars show average± SD; unpaired Student’s t-tests were performed, *P<0,05. D)- HCT116 EGFP-LC3 TetOn-M2 WT and ATG4D -/- clone 25 cells stable expressing mCherry-ATG4D or mCherry alone, were treated for 8 hours with 10 μM doxycycline and analysed by western blotting. E)- Quantification of D. The graph shows the change in LC3II/LC3I ratio from 3 independent experiments; bars show average± SD; unpaired Student’s t-tests were performed, **P < 0.01, *P<0,05.

The different isoforms of ATG4 have been reported to exhibit some specificity for individual ATG8 paralogues and it has been suggested that the GABARAP subfamily is particularly susceptible to changes in de-lipi-dation conditions (Kauffman et al., 2018). We observed similar increases in levels of lipidation of GABARAP-L1 and GABARAP-L2 in ATG4D −/− cells (supplementary figure 5B).

To further validate the effect of ATG4D depletion on LC3-lipidation, we performed rescue experiments. In ATG4D −/− clone 25, M2-induced LC3B lipidation levels are rescued upon expression of ATG4D-mCherry, but not in cells expressing the mCherry control plasmid (figure 5D and E). We therefore conclude that ATG4D plays a role in the de-lipidation of LC3/GABARP-like molecules during non-canonical autophagy.

## Discussion

The subtractive CRISPR screen described here correctly identified ATG7, ATG10, ATG3 and the ATG16L1-ATG5-ATG12 complex, all of which are known to be enzymatically required for LC3-lipidation in any context. In contrast, sgRNAs targeting genes such as FAM208A and MAX, both required for expression of the EGFP-LC3 transgene, provide strong signals on both individual screens but are not detected in the subtraction results (Supplementary figure 1, compare A and B). All sgRNAs targeting these genes lie close to the y=x line on a z-z plot (figure 2B), demonstrating the specificity of the subtraction approach.

Although our screen was not optimised to detect genes essential for cell viability, we nonetheless identified a v-ATPase subunit. We previously identified the WD40 CTD of ATG16L1 as required for LC3-lipidation in response to M2-proton channel activity, ionophores and during LAP (Fletcher et al., 2018). The recently described vATPase/ATG16L1 interaction identified during salmonella infection also depends on this domain (Xu et al., 2019). This suggested that M2-and ionophore-induced LC3-lipidation might depend on the same mechanism. The antagonism of these processes by the salmonella effector SopF and the concomitant association of ATG16L1 and V-ATPase suggest this is the predominant mechanism by which non-canonical lipidation of LC3 and other ATG8-like molecules occurs. The v-ATPase consists of many subunits that assemble into the transmembrane VO subcomplex and the cytosolic V1 subcomplex. The reversible association and dissociation of these subcomplexes regulates the activity of the v-ATPase, where the associated form is active (Kane, 1995; Oot et al., 2017; Sumner et al., 1995). Interestingly, BafA1, an inhibitor of WD40 CTD-dependent LC3-lipidation, promotes v-ATPase disassembly (Poëa-Guyon et al., 2013). LC3-lipidation in response to ionophores and during entosis can be inhibited with the v-ATPase inhibitor bafilomycin A1 (Fletcher et al., 2018; Florey et al., 2015). Further experiments will be required to define a precise role for v-ATPase assembly state in ATG16L1 recruitment.

We identified the RalGAP complex as important for M2-induced LC3-lipidation. We were able to demonstrate that lack of either RalGAPβ alone or RalGAPα1 and RalGAPα2 expression inhibits LC3-lipidation caused by M2 expression. In the context of IAV infection the observed reduction in LC3-lipidation was accompanied by a modest reduction in M2 expression levels, though this did not substantially impair viral replication. However, no phenotype was observed in the other WD40 CTD-dependent LC3-lipidation pathways indicating that an effect of RalGAP signaling could be M2 specific. These data are consistent with an effect of RalGAP on M2, for instance its sub-cellular localisation, that affects LC3-lipidation indirectly. The subtractive screen identified ATG4D as a negative regulator of LC3-lipidation, and ATG4D knock out indeed resulted in increases levels of lipidated LC3B and GABARAPL1 and L2. The contributions of the individual ATG4 paralogues to priming and delipidation of ATG8s are still not fully understood. Paralogues appear to exhibit a preference for priming or delipidation of individual ATG8s. However, other paralogues appear to be able to compensate for loss of individual paralogues to some degree (Agrotis et al., 2019; Kauffman et al., 2018). ATG4D is the least well characterised paralogue, in part due to low activity in vitro (Kauffman et al., 2018). In our system, ablation of ATG4D expression resulted in increased LC3B-II in the context of M2-induced LC3-lipidation, as well as under basal conditions. Similar observations were made for GABAR-AP-L1 and GABARAP-L2. Elsewhere we describe a role for ATG4D in delipidation of phosphatidylserine conjugated LC3, which accumulates during WD40 CTD-dependent LC3-lipidation – including in response to IAV M2 – but not during canonical autophagy (Durgan et al., 2020). A missense mutation in ATG4D was found in Lagotto Romagnolo dogs with progressive neurological symptoms (Kyöstilä et al., 2015). These dogs exhibited increased levels of LC3-II in basal and starvation induced autophagy (Syrjä et al., 2017). Whether the neurological phenotype is linked to reduced recycling of phosphatidylserine conjugated LC3 remains to be determined.

Our data are consistent with a simple model in which recruitment of ATG16L1 by the v-ATPase targets lipidation of LC3 to erroneously neutral compartments. Targeting of this pathway by diverse pathogens such as salmonella and influenza suggests it may represent an important damage detection mechanism, as even minimal damage to a compartment would compromise the ability of the v-ATPase to maintain proton gradients.

## Materials and Methods

### Antibodies and Reagents

Antibodies used in this study are anti-LC3B (NB100-2220 and NBP2-46892, Novus Biologicals, WB 1:1000 and 1: 2000 respectively), anti-Influenza A Virus M2 Protein antibody [14C2] (ab5416, Abcam, WB 1:1000, IF: 1:100, FACS 1:100 and GTX125951, Genetex, WB 1:1000), Anti-Influenza A Virus Nucleoprotein (ab20343, Abcam, WB 1:2000 and GTX125989, Genetex, WB 1:5000), anti-GABARP-L1 (11010-1-AP, Proteintech, WB 1:1000), anti-GABARP-L2 (ab126607, Abcam, WB 1:1000), anti-RalGAPα1 (ab182570, Abcam, WB 1:10,000), anti-RalGAPβ (ab151139, Abcam, WB 1:1000), anti-ATG4D (16924-1-AP, Proteintech, WB 1:1000), anti-mCherry (NBP1-96752, Novus Biologicals, WB 1:1000),), anti-ATG16L1 (8089, Cell Signaling Technology, WB 1:1000), an-ti-ATG16L1 (PM040, MBL International Corporation, IP), anti-ATG12 (sc271688, Santa Cruz, WB 1:1000), anti-ATP6V1D (ab157458, Abcam, IP and WB 1:1000), anti-ATP6V1A (ab199326, Abcam, WB 1:1000), anti-ATP6V1B1/2 (sc55544, Santa Cruz, WB 1:1000), anti-FLAG (F1804, Sigma-Aldrich, IP and WB 1:1000), anti-β-Actin (20536-1-AP and 66009-1, Proteintech, WB 1:10000), anti-GAPDH (ab8245, Abcam, WB 1:10000) and anti-tubulin (AbD serotec MCA77G, WB 1:2000). Torin 1 (S2827) and VPS34 inhibitor 1 (Compound 19, PIK-III analogue, S8456) were obtained from Selleckchem. Doxycycline (D9891), monensin (M5273), Wortmannin (W1628) and amantadine hydrochloride (A1260) were obtained from Sigma. Bafilomycin A1 (ab120497) was purchased from Abcam. Polybrene (sc134220) was obtained from Santa Cruz.

### Cell culture and RNA interference

HCT116 cells were cultured in McCoys 5A medium (Lonza) supplemented with 5% foetal calf serum (FCS). HEK293t and MDCK-II (a kind gift from P. Digard, Roslin Institute, Edinburgh, UK) were cultured in Dulbecco’s modified Eagle’s medium (DMEM; Gibco Life Technologies) containing 10% FCS. HCT116 EGFP-LC3B TetON M2 expresses the IAV strain Udorn M2 protein under a doxycycline inducible promoter. It was generated by transducing HCT116 EGFP-LC3B cells using a lentivirus produced with pInd10b-Ud-M2 followed by initial selection with G418. A high expressing cell clone was isolated using FACS. All cells were maintained in an incubator at 37°C with 5% CO2. For siRNA experiments, cells were treated with 40 nM of non-targeting (NT1) or target-specific siRNA oligonucleotides (Dharmacon On-Target Plus Smart Pool), using Lipofectamine RNAi-MAX (Invitrogen) according to manufacturer’s instructions.

### siRNAs and Plasmids

Sequences of siRNAs used in this manuscript were as follows: ATG4D smart pool (5’-CGGACCAGCUUUAGCAAGA-3’, 5’-GAAGGCAGGU-GACUGGUAU-3’, 5’-CAAGUACGGUUGGGUGGUU-3’, 5’-AGGGU-GACAUACAGCGUUU-3’).

M4P-EGFP-LC3B and pOPG was a kind gift from F. Randow. pMD2.G (Addgene plasmid # 12259; http://n2t.net/addgene:12259; RRID:Ad-dgene_12259) and psPAX2 (Addgene plasmid # 12260; http://n2t.net/ addgene:12260; RRID:Addgene_12260) were a gift from Didier Trono. pSpCas9(BB)-2A-Puro (PX459) V2.0 was a gift from Feng Zhang (Addgene plasmid # 62988; http://n2t.net/addgene:62988; RRID:Ad-dgene_62988) (Ran et al., 2013).

The insert encoding the M2 protein from IAV virus strain Udorn was generated by overlap extension PCR using pDual-seg7-Ud (a kind gift from P. Digard) as a template and cloned into pInd10b-HA-KRAS (L. Lee, unpublished) using AgeI and MluI sites by standard restriction ligation cloning.

M5P-mCherry-SopF was subcloned into M5PmCherry-hATG16L1 (R. Ulferts, unpublished) from M4P-EGFP-SopF (a kind gift from F. Randow) using the PciI and NotI restriction sites.

M5P-mCherry control vector was generated by inserting a stop codon downstream of mCherry by inserting a linker generated by annealing 5’-CATGTCGTAAGTAATTAAGC-3’ and 5’-GGCCGCTTAATTACT-TACGA-3’ into the PciI and NotI restriction sites of M5P-mCherry-hATG16L1.

pBABE-FLAG-S-mATG16L1 was described in Gammoh et al., 2013)(a kind gift from N. Gammoh).

plenti-ATG4D-mCherry-hygR was generated by gateway cloning with LR clonase (ThermoFisher Scientific) using pENTR-ATG4D (Transomics) and pLenti-GWT-mCherry-HygR (a kind gift from F. Sorgeloos). pLenti-PGK-RalGAPA1-hygB, pLenti-CMV-RalGAPA1-hygB, and pLenti-mCherry-RalGAPA1 were generated by gateway cloning with LR clonase (ThermoFisher Scientific) using pENTR-RalGAPA1 (Transomics) and, pLenti-PGK-GWT-hygB (a kind gift from F. Sorgeloos), pLenti CMV hygro DEST (a gift from Eric Campeau & Paul Kaufman (Addgene plasmid # 17454; http://n2t.net/addgene:17454; RRID:Addgene_17454) or pLenti-mCherry-GWT (a kind gift from F. Sorgeloos).

### Whole genome CRISPR screen

Human GeCKOv2 CRISPR knockout pooled library and lenti-Cas9-Blast was a gift from Feng Zhang (Addgene # 1000000049, (Sanjana et al., 2014)). The library was amplified and the lentivirus library generated as described in (Shalem et al., 2014). HCT-116 stably expressing EGFP-LC3B tet-ON M2 were transduced with cas9-blast lentivirus and selected with blasticidin. Cells were transduced with the virus library at a 300x library representation at an moi of 0.3 viruses per cell, followed by selection with puromycin. Selected cells were passaged at 300x library representation for 14 days. Expression of M2 was induced by addition of doxycycline. 16 h post induction cells were harvested and stained for M2 surface expression using anti-M2 antibody (14C2) followed by staining with donkey-anti-mouse-IgG-Alexa568 (Thermo). Cells were then permeabilised with 0.1% saponin (Sigma) in PBS or mock treated. Zombie-violet (Biolegend, 423114, 1:200) staining was used to exclude dead cells. Cells were fixed with 1% formaldehyde in PBS prior to sorting. The top and bottom 10% EGFP-expressing cells of the M2-positive cell populations were collected on a BD FACSAriaTM Fusion or BD FACSJazzTM instrument. Genomic DNA was isolated using the QIAamp DNA FFPE Tissue Kit (Qiagen) essentially as described in the manufacturers protocol except that buffer ATL was supplemented with 300 mM NaCl and the cell lysate incubated for 2 h at 56°C under constant agitation. PCR amplification was carried out as described in (Shalem et al., 2014). Illumina Hiseq was performed at the Bauer sequencing facility, Harvard.

### Data analysis of the CRISPR screen

For analysis of the screen data, we devised a simple but robust subtraction approach based on our previous method for genome-wide screen analysis (Li et al., 2020). Briefly, read counts corresponding to each guide RNA were normalised to reads per million and and log transformed. Quantile normalisation was performed in R version 3.6.1. In comparisons between intervention and control experiments, over/under-representation was quantified as the distance from the expected null (i.e. the y=x line on a plot of read counts.) In order to control for heteroscedasticity, these distances were normalised to local z-scores calculated for sliding bins of adjacent read count results (Li et al., 2020).

In order to remove the background effects of specific genes required for expression of the EGFP construct, z-scores from the background (EGFP expression) screen were then subtracted from z-scores for the saponin-permeabilisation (M2) screen. p-values were calculated from the sum of z-scores for sgRNAs targeting a particular gene compared to a density function modelled on an empirical distribution of possible combinations of sgRNA z-scores permuted at least 1e8 times by randomly rearranging z-scores for all sgRNAs in the screen.

### CRISPR knock out of single genes

Stable knock out cell lines using CRISPR technology were generated using either plasmid or nucleofection with guide RNAs (Synthego) and Cas9 (Thermo). Single cell clones were selected and absence of gene expression confirmed by western blotting or qPCR.

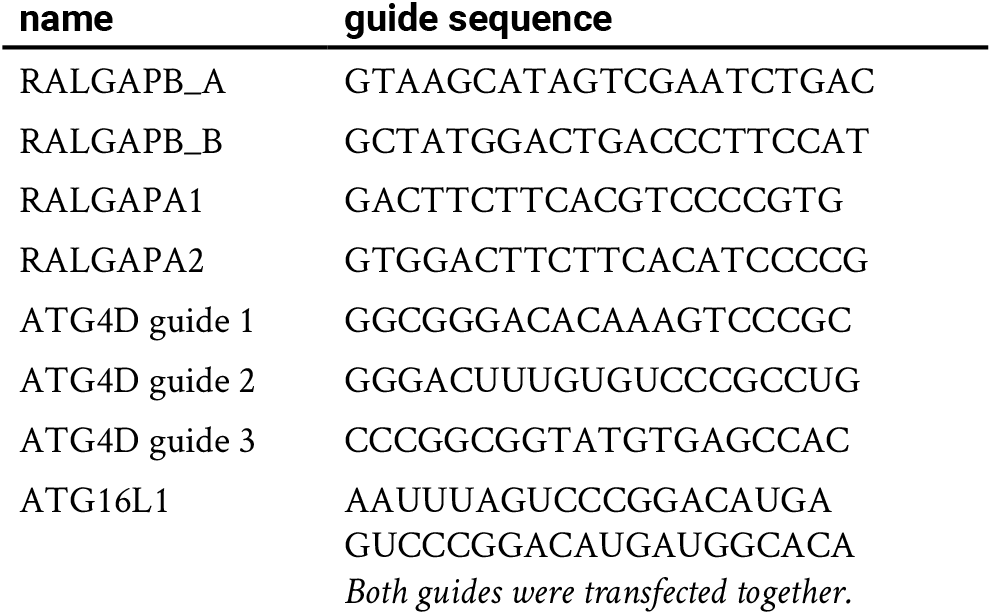

### qPCR

RNA was isolated using RNAeasy extraction Kit Qiagen following manufacturer instructions. cDNA was synthesised using SuperSCRIPT-II reverse transcriptase and random hexamer primer according to manufactures protocol. qPCR was performed using taq PCR and cycler using primers GCCTGGATAACCAGTCTTCTCC and CACAGATCAGCCTGTAGGCTTG for RalGAPα2 and GGGGTGTTGAAGGTCTCAAA and TTCTACAATGAGCTGCGTGTG for actin.

### Influenza A Virus production

Stocks of influenza A virus PR8 (strain A/Puerto Rico/8/1934) and MUd, a reassortant PR8 variant carrying segment 7 of IAV strain A/ Udorn/307/1972 (Noton et al., 2007) were generated using the eight plasmid-based systems as previously described (Wit et al., 2004) and propagated on MDCK-II cells in presence of TPCK-trypsin (Worthington). For infection, cells were first washed with serum free medium and incubated with virus in serum-free medium at 37°C. After 1 h, the medium was replaced with DMEM containing 1% FCS. Virus titres were determined by plaque assay on MDCK-II cells.

### Retrovirus and lentivirus production

Retrovirus and lentivirus particles were generated using packaging plasmids MD2-G and pOPG or psPAA2, respectively, by transfecting HEK-293T using PEI. Cells were transduced with virus by spinfection in the presence of 8 μg/ml of polybrene followed by selection with G418, blasticidin, puromycin or fluorescence-assisted cell sorting as appropriate.

### Entosis assay

The entosis assay was carried out as described previously (Florey et al., 2011). HCT116 wild type or knock out cells stably expressing EGFP-LC3B were grown in 35 mm glass bottom dishes and imaged every 4 min for 20 h. DIC and fluorescent images were acquired using a confocal Zeiss LSM 780 microscope (Carl Zeiss Ltd) equipped with a 40x oil immersion 1.40 numerical aperture (NA) objective using Zen software (Carl Zeiss Ltd).

### Western blotting

Cells were lyses in ice cold RIPA (10 mM Tris-HCl pH 7.5, 150 mM NaCl, 1% Triton X-100, 0.1% SDS, 1% sodium deoxycholate) or NP40 buffer (0.5% NP-40, 25 mM Tris-HCl pH 7.5, 100 mM NaCl, 50 mM NaF) supplemented with cOmplete^TM^, Mini, EDTA-free Protease Inhibitor Cocktail (11836170001). Lysates were cleared by centrifugation and the protein concentration determined using BCA assay (Pierce) and IgG or BSA as a standard. Proteins were separated in a Mini-PROTEAN®TGX™ gel (BioRad) and transferred onto nitrocellulose. After blocking with 5 or 10% dry milk powder in TBS supplemented with 0.1% Tween 20, blots were incubated for 1 h to overnight with primary antibody at the indicated dilution, followed by the appropriate species specific IRdye 800CW and 680LT coupled secondary antibodies (LICOR) and imaged using an Odyssey CLx scanner (Li-COR). Analysis and quantification were carried out using Imagestudio (Li-COR). In some cases, antibodies were subsequently removed using Restore Stripping buffer (ThermoFisher Scientific; #21059) according to manufacturer’s instructions, reblocked and incubated with primary and secondary antibody.

### Immunofluorescence

Cells were grown on coverslips pre-treated with 0,001% Poly-L-Lysine (Sigma), fixed using 4% formaldehyde in PBS, permeabilised with 0,05% saponin and incubated with primary antibody, prior to staining with AlexaFluor 405, 488, 568 or 647 coupled secondary antibodies. Images were acquired using Zeiss LSM800 with Airyscan and further processed using Adobe Photoshop CC2020 or Fiji 1.0 (Rueden et al., 2017; Schindelin et al., 2012).

### Immunoprecipitation (IP)

This protocol was adapted from Xu et al., 2019. Two 15cm dishes of cells were washed twice with ice-cold PBS and lysed in a buffer containing 50 mM Tris-HCl (pH 7.5), 150 mM NaCl, 2 mM EDTA, 0.8% C12E9 (Sigma-Aldrich, P9641) and protease inhibitors (Roche, 11836170001 or Sig-ma-Aldrich, P8340). Lysates were centrifugated at 13,000 rpm, 4°C for 30 minutes and the pellet discarded. A small amount of lysate was removed for western blotting. 60 μL of Dynabeads® Protein A or Protein G (Invitrogen, 10002D and 10004D) were incubated with 5 μL of IP antibody for 30 minutes rotating at 4°C. Beads were washed once in lysis buffer before incubation with lysate for 2h rotating at 4°C. Beads were then washed five times in a buffer containing 50 mM Tris-HCl (pH 7.5), 150 mM NaCl, 2 mM EDTA, 1% Triton X-100, and 0.1% C12E9. Bound proteins were eluted by boiling at 95°C for 5 minutes in SDS-loading buffer. Eluted samples were probed by western blot as described above.

## Acknowledgments

We thank Sharon Tooze, Noor Gammoh, Frederic Sorgeloos, Paul Lehner, Paul Digard, Felix Randow and Keith Boyle for helpful discussions and provision of reagents. This work was funded by the Medical Research Council (MR/M00869X/1, R.B.) and supported by the Cambridge NIHR BRC Cell Phenotyping Hub, the Francis Crick Institute Science Technology Platforms Advanced Light Microscopy, Flow Cytometry and Cell Services.

## Author contributions

R.U. and E.M conceptualized the study and designed the experiments. R.U., L.T., E.M., and R.B. wrote the paper. R.U. and R.B. designed the CRISPR screen. J.K.B. designed and performed the data analysis of the screen. R.U., E.M., L.T., B.M. and O.F. performed the experiments. A.D., L.C.L. and S.D.T. provided crucial reagents.

---

**Figure S. 1.**
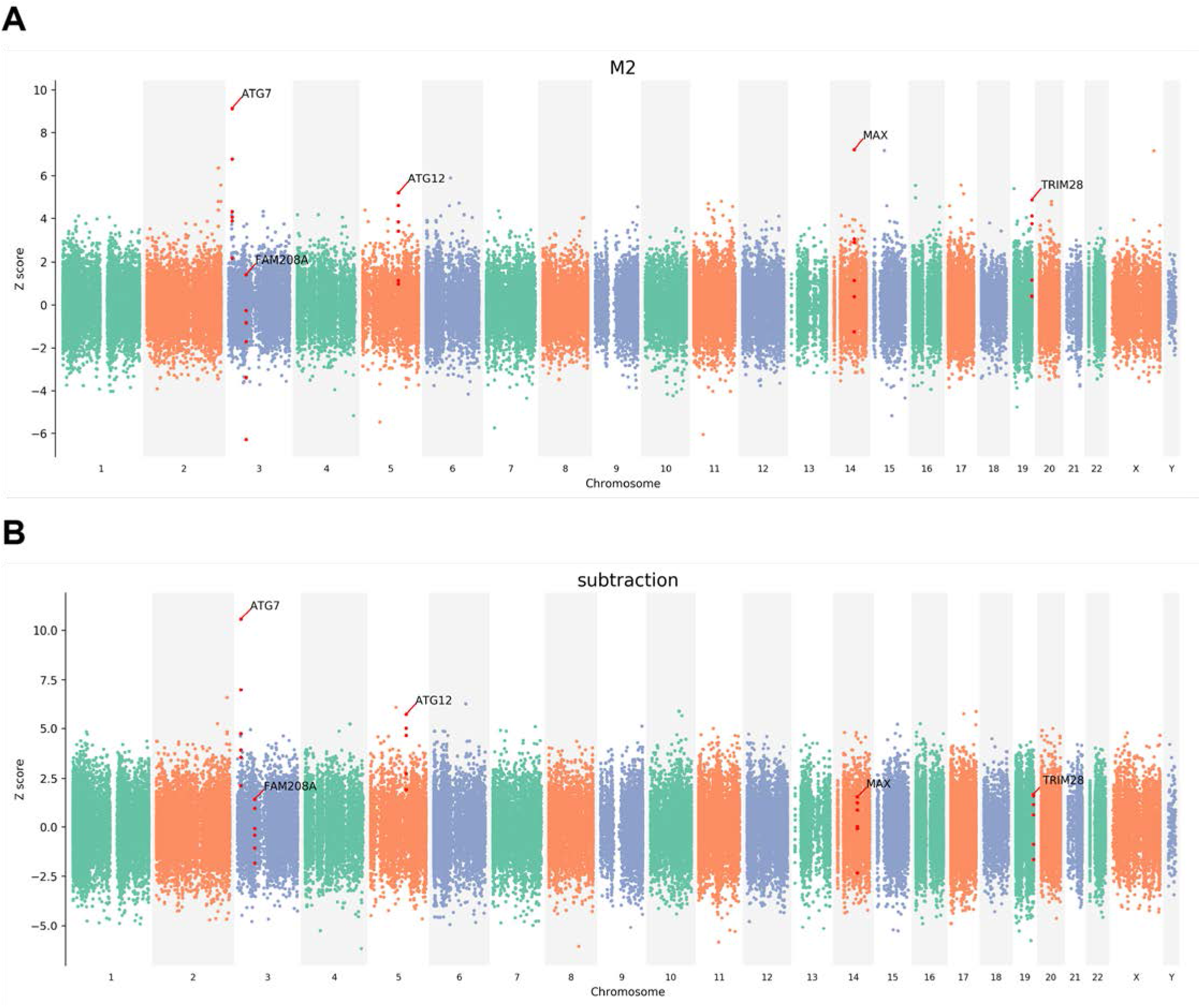
Spatial distribution on the genome of the z-score of sgRNAs the A) permeabilised FACS sort of EGFP-LC3B levels and B) the substractive analysis of permeabilised and unpermeabilised FACS sorts.?

**Figure S. 2.**
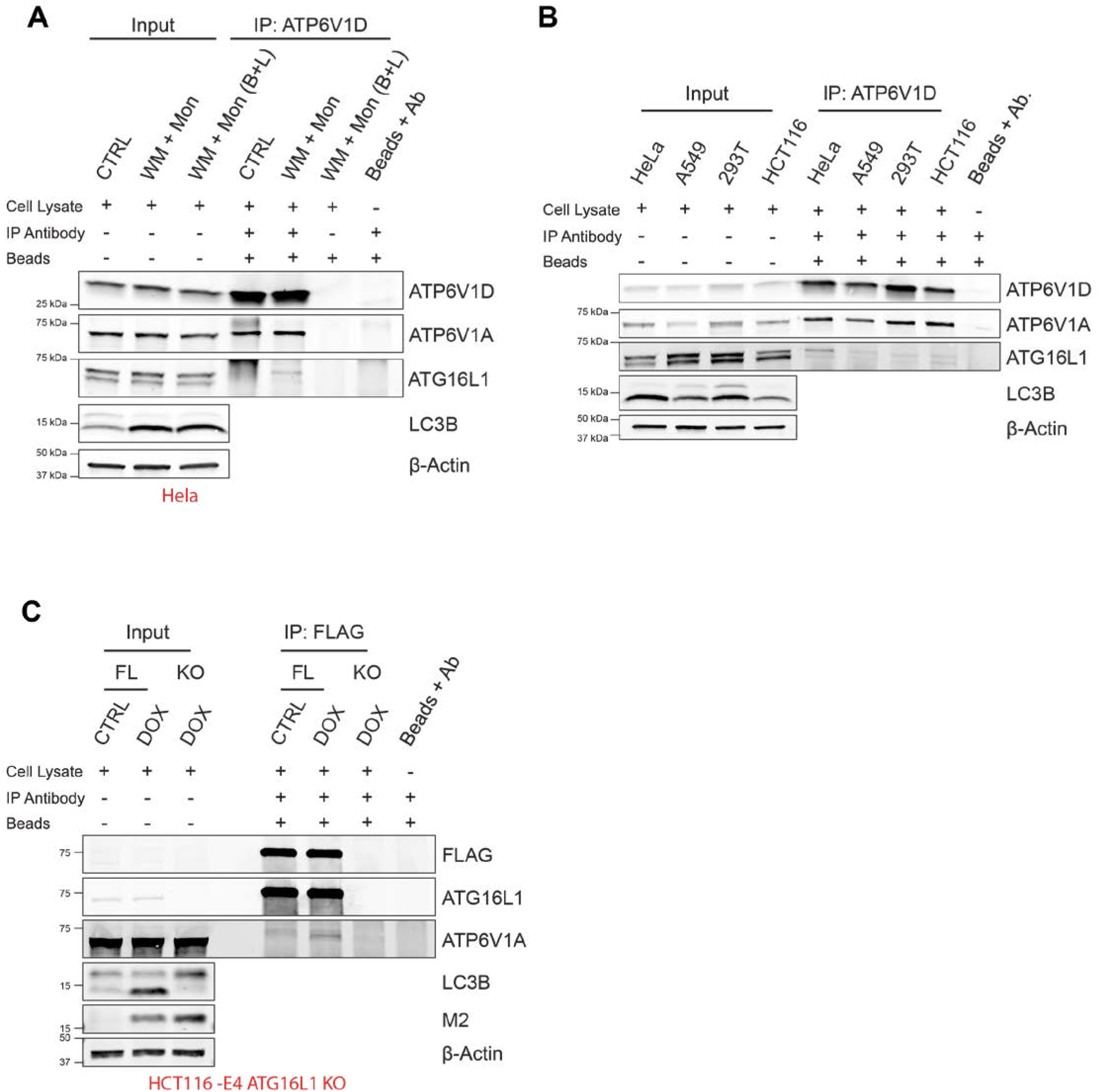
A) Immunoprecipitation analysis of endogenous of ATP6V1D in HeLa cells treated with wortmannin (IN-1 substitute; pretreatment: 100 nM for 30 minutes) followed by monensin (100 μM for 1h). B) Immunoprecipitation analysis of endogenous of ATP6V1D in the indicated cell lines following treatment with VPS34 IN-1 (pretreatment: 1 μM for 30 minutes) followed by monensin (100 μM for 1h). C) Immunoprecipitation analysis of FLAG-S-mATG16L1 in HCT116 EGFP-LC3B TetON-M2 ATG16L1 −/− cells following doxycycline treatment (10 μg/mL for 16h). FL: FLAG-S-mATG16L1 transduced; KO: untransduced.

**Figure S. 3.**
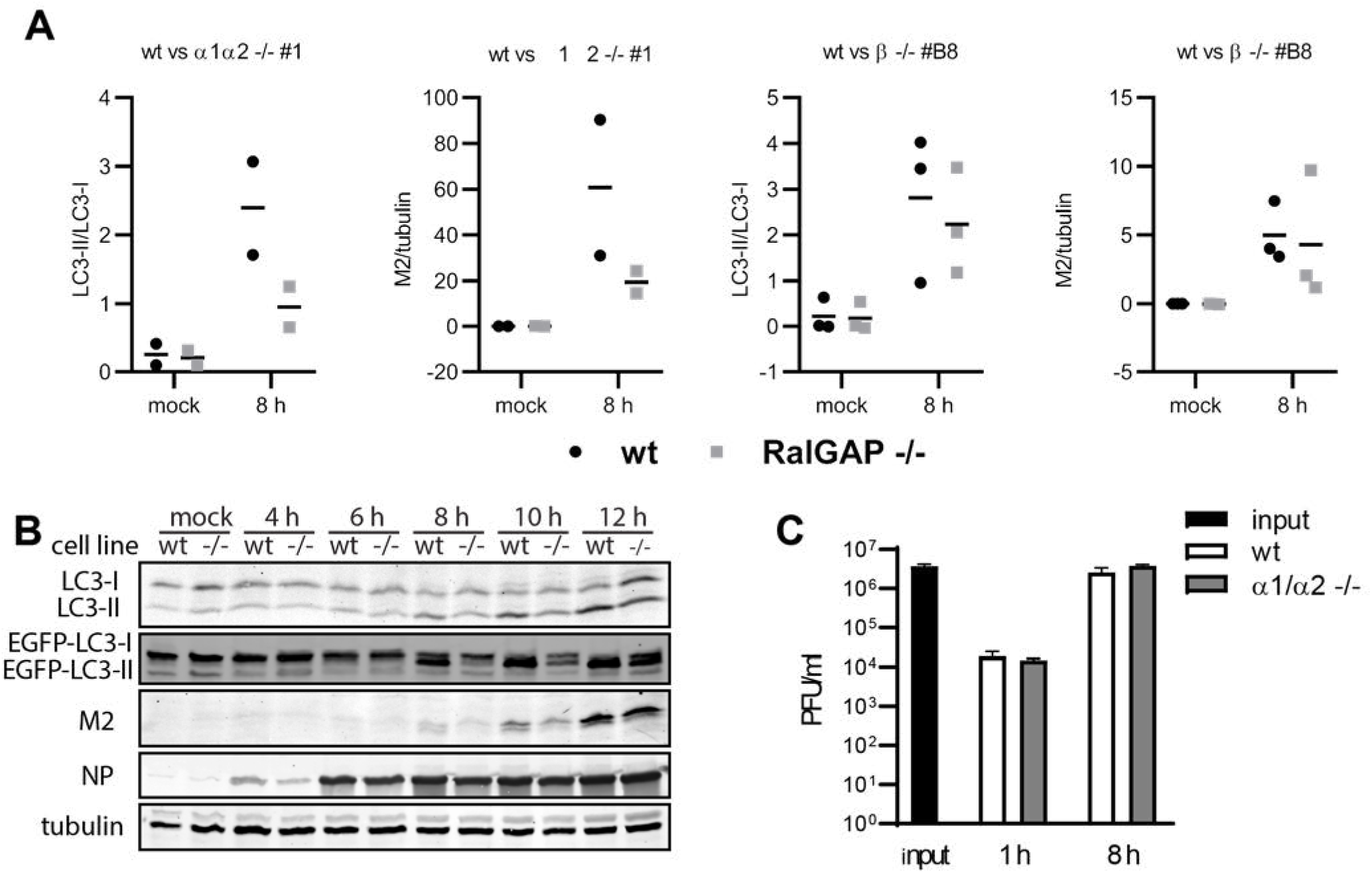
A) Quantification of LC3-lipidation analysis by western blot of HCT116 EGFP-LC3B TetON-M2 cells depleted for RalGAPα1α2 (panel 1 and 2) or RalGAPβ (clone B8) after infection with PR8 at an MOI of 10 PFU per cell for 8 h. Graphs show mean of LC3II/LC3I ratio (panels 1 and 3) and M2 expression normalised to tubulin (panels 2 and 4) from two or three independent experiments. B) LC3-lipidation analysis of HCT116 EGFP-LC3B TetON-M2 wt and cells depleted for RalGAPα1α2 infected with MUd at an MOI of 10 PFU per cell and lysed at the indicated time point p.i.. C) HCT116 EGFP-LC3B TetON-M2 wt or cells depleted for RalGAPα1α2 were infected at an MOI of 10 PFU per cell. Supernatants were harvested at 1 or 8 h p.i. and titres determined by plaque assay.

**Figure S. 4.**
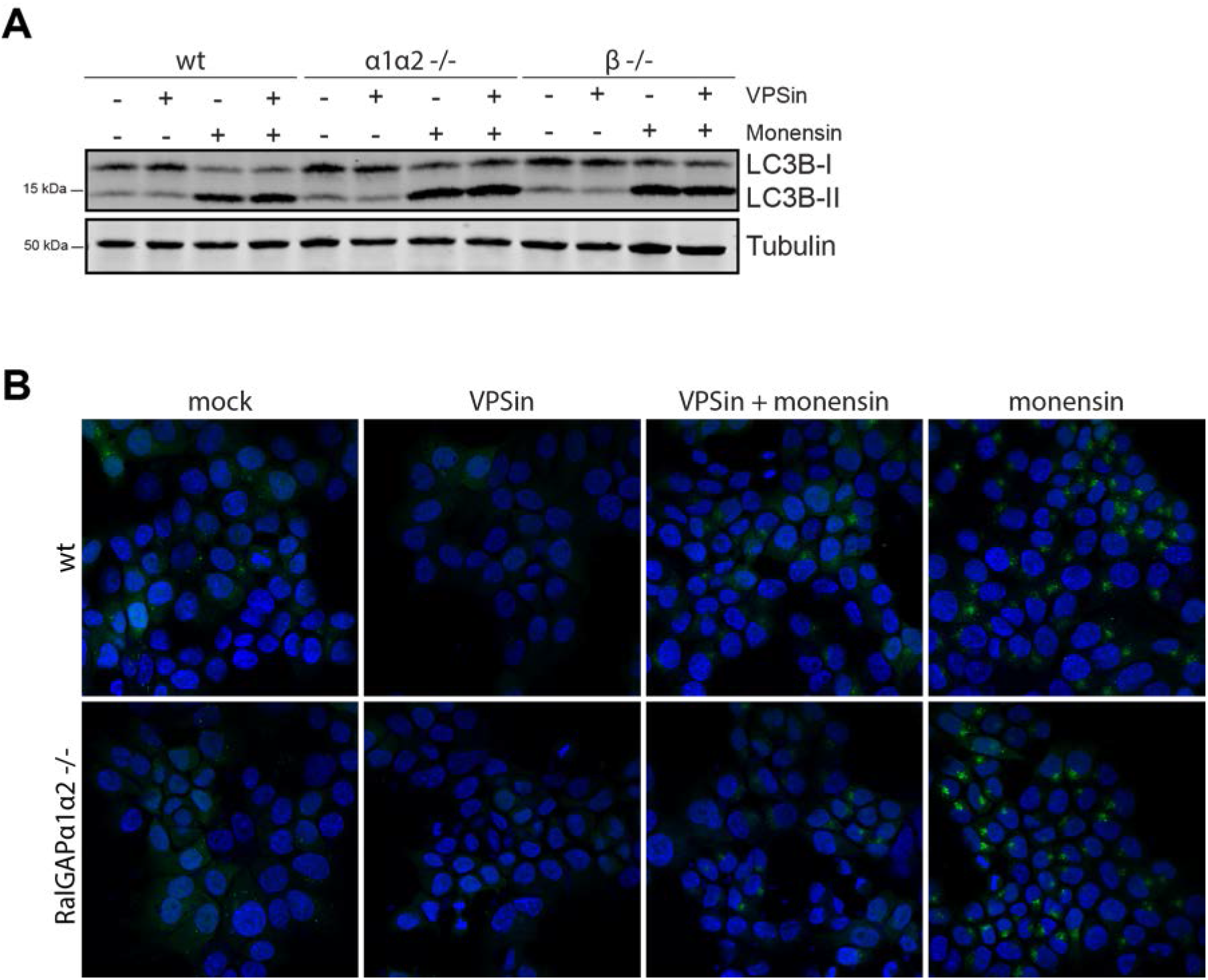
A) LC3 lipidation analysis of HCT116 EFGP-LC3B TetON M2 wt and RalGAPα1α2−/− cells after 30 min treatment with 1 μM monensin. Where indicated cells were pretreated with 1 μM VPSin for 20 min prior to addition of 1 μM monensin to inhibit canonical LC3-lipidation. B) Representative images of IF analysis of HCT116 EFGP-LC3B TetON M2 wt and RalGAPα1α2−/− treated as in B.

**Figure S. 5.**
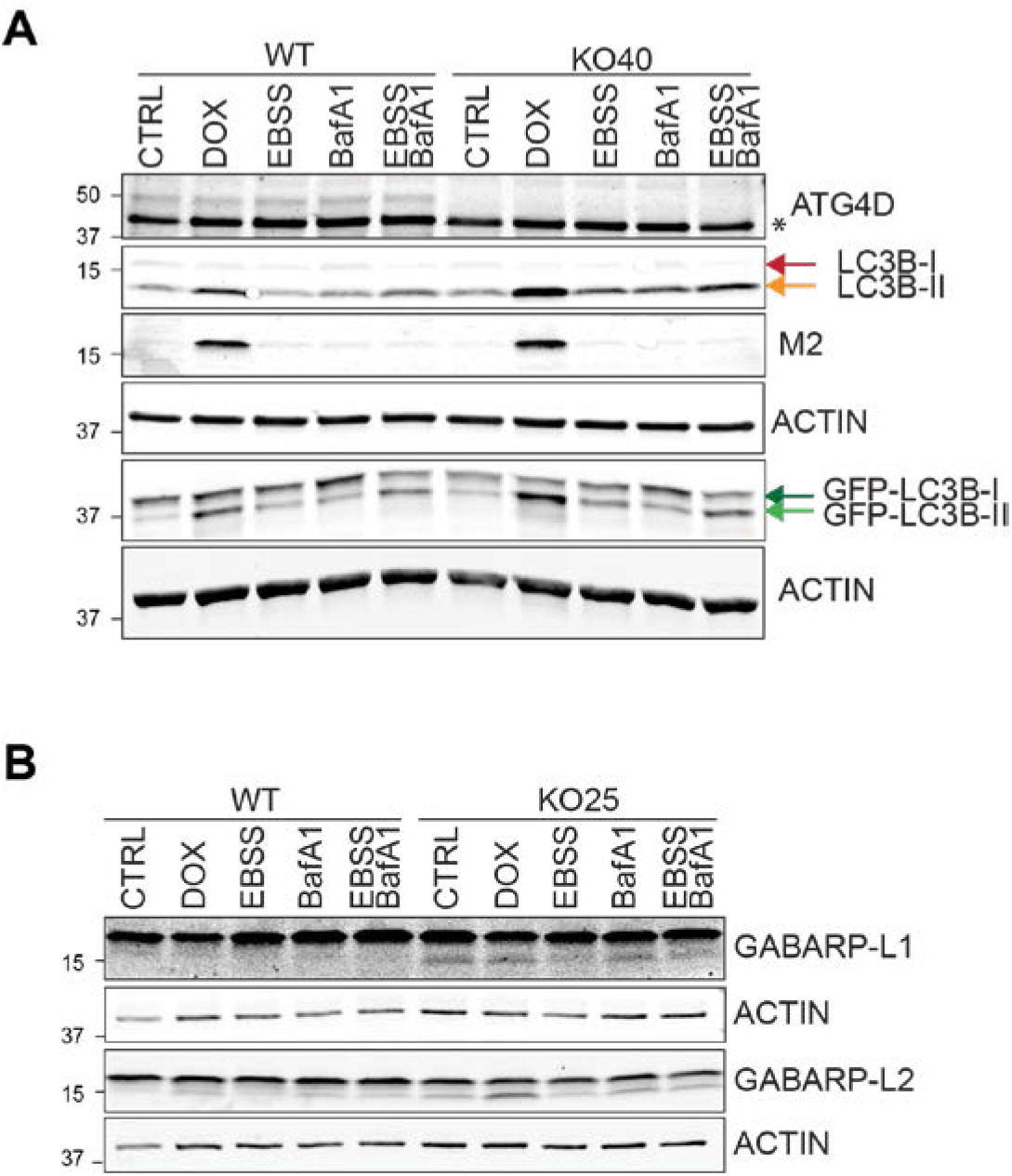
A) HCT116 EGFP-LC3 TetOn-M2 WT and ATG4D KO clone 40 were treated with 10 μg/ml dox for 8 h, 2 hr with EBSS, 1 hr with 200 nM Bafilomycin A1 and a combination of EBSS together with Bafilomycin A1. arrowhead indicates ATG4D specific band, * background band. B) HCT116 EGFP-LC3 TetOn-M2 WT and ATG4D -/clone 25 were treated as in B and GABARAP-L1 and GABARAP-L2-lipidation analysed by western blot.

## References

Agrotis, A., Pengo, N., Burden, J.J., and Ketteler, R. (2019). Redundancy of human ATG4 protease isoforms in autophagy and LC3/GABARAP processing revealed in cells. Autophagy 15, 976–997.

Beale, R., Wise, H., Stuart, A., Ravenhill, B.J., Digard, P., and Randow, F. (2014). A LC3-Interacting Motif in the Influenza A Virus M2 Protein Is Required to Subvert Autophagy and Maintain Virion Stability. Cell Host & Microbe 15, 239–247.

Ciampor, F., Bayley, P.M., Nermut, M.V., Hirst, E.M.A., Sugrue, R.J., and Hay, A.J. (1992). Evidence that the amantadine-induced, M2-mediated conversion of influenza A virus hemagglutinin to the low pH conformation occurs in an acidic trans golgi compartment. Virology 188, 14–24.

Durgan, J., Lystad, A.H., Sloan, K., Carlsson, S.R., Wilson, M.I., Marcassa, E., Ulferts, R., Webster, J., Lopez-Clavijo, A.F., Wakelam, M.J., et al.(2020). Non-canonical autophagy drives alternative ATG8 conjugation to phosphatidylserine. BioRxiv 2020.05.14.096115.

Eng, K.E., Panas, M.D., Hedestam, G.B.K., and McInerney, G.M. (2010). A novel quantitative flow cytometry-based assay for autophagy. Autophagy 6, 634–641.

Fletcher, K., Ulferts, R., Jacquin, E., Veith, T., Gammoh, N., Arasteh, J.M., Mayer, U., Carding, S. R., Wileman, T., Beale, R., et al. (2018). The WD40 domain of ATG16L1 is required for its non-canonical role in lipidation of LC3 at single membranes. The EMBO Journal 37, e97840.

Florey, O., Kim, S.E., Sandoval, C.P., Haynes, C.M., and Overholtzer, M. (2011). Autophagy machinery mediates macroendocytic processing and entotic cell death by targeting single membranes. Nat Cell Biol 13, 1335–1343.

Florey, O., Gammoh, N., Kim, S.E., Jiang, X., and Overholtzer, M. (2015). V-ATPase and osmotic imbalances activate endolysosomal LC3 lipidation. Autophagy 11, 88–99.

Fujioka, Y., Suzuki, S.W., Yamamoto, H., Kondo-Kakuta, C., Kimura, Y., Hirano, H., Akada, R., Inagaki, F., Ohsumi, Y., and Noda, N.N. (2014). Structural basis of starvation-induced assembly of the autophagy initiation complex. Nat Struct Mol Biol 21, 513–521.

Gammoh, N., Florey, O., Overholtzer, M., and Jiang, X. (2013). Interaction between FIP200 and ATG16L1 distinguishes ULK1 complex-dependent and -independent autophagy. Nat Struct Mol Biol 20, 144–149.

Gannagé, M., Dormann, D., Albrecht, R., Dengjel, J., Torossi, T., Rämer, P.C., Lee, M., Strowig, T., Arrey, F., Conenello, G., et al. (2009). Matrix Protein 2 of Influenza A Virus Blocks Autophagosome Fusion with Lysosomes. Cell Host & Microbe 6, 367–380.

Henkel, J.R., Popovich, J.L., Gibson, G.A., Watkins, S.C., and Weisz, O.A. (1999). Selective Perturbation of Early Endosome and/ortrans-Golgi Network pH but Not Lysosome pH by Dose-dependent Expression of Influenza M2 Protein. Journal of Biological Chemistry 274, 9854–9860.

Jacquin, E., Leclerc-Mercier, S., Judon, C., Blanchard, E., Fraitag, S., and Florey, O. (2017). Pharmacological modulators of autophagy activate a parallel noncanonical pathway driving unconventional LC3 lipidation. Autophagy 0, 1–14.

Kane, P.M. (1995). Disassembly and Reassembly of the Yeast Vacuolar H+-ATPase in Vivo. J. Biol. Chem. 270, 17025–17032.

Kauffman, K., Yu, S., Jin, J., Mugo, B., Nguyen, N., O’Brien, A., Nag, S., Lystad, A., and Melia, T. (2018). Delipidation of mammalian Atg8-family proteins by each of the four ATG4 proteases. Autophagy 14, 992–1010.

Kaufmann, A., Beier, V., Franquelim, H.G., and Wollert, T. (2014). Molecular Mechanism of Autophagic Membrane-Scaffold Assembly and Disassembly. Cell 156, 469–481.

Kyöstilä, K., Syrjä, P., Jagannathan, V., Chandrasekar, G., Jokinen, T.S., Seppälä, E.H., Becker, D., Drögemüller, M., Dietschi, E., Drögemüller, C., et al. (2015). A Missense Change in the ATG4D Gene Links Aberrant Autophagy to a Neurodegenerative Vacuolar Storage Disease. PLOS Genetics 11, e1005169.

Li, B., Clohisey, S.M., Chia, B.S., Wang, B., Cui, A., Eisenhaure, T., Schweitzer, L.D., Hoover, P., Parkinson, N.J., Nachshon, A., et al. (2020). Genome-wide CRISPR screen identifies host dependency factors for influenza A virus infection. Nature Communications 11, 1–18.

Martin, T.D., Chen, X.-W., Kaplan, R.E.W., Saltiel, A.R., Walker, C.L., Reiner, D.J., and Der, C.J. (2014). Ral and Rheb GTPase Activating Proteins Integrate mTOR and GTPase Signaling in Aging, Autophagy, and Tumor Cell Invasion. Molecular Cell 53, 209–220.

Mizushima, N., and Komatsu, M. (2011). Autophagy: Renovation of Cells and Tissues. Cell 147, 728–741.

Mizushima, N., Noda, T., Yoshimori, T., Tanaka, Y., Ishii, T., George, M.D., Klionsky, D.J., Ohsumi, M., and Ohsumi, Y. (1998). A protein conjugation system essential for autophagy. Nature 395, 395–398.

Noton, S.L., Medcalf, E., Fisher, D., Mullin, A.E., Elton, D., and Digard, P. (2007). Identification of the domains of the influenza A virus M1 matrix protein required for NP binding, oligomerization and incorporation into virions. Journal of General Virology, 88, 2280–2290.

Oot, R.A., Couoh-Cardel, S., Sharma, S., Stam, N.J., and Wilkens, S. (2017). Breaking up and making up: The secret life of the vacuolar H+-ATPase. Protein Science 26, 896–909.

Poëa-Guyon, S., Ammar, M.R., Erard, M., Amar, M., Moreau, A.W., Fossier, P., Gleize, V., Vitale, N., and Morel, N. (2013). The V-ATPase membrane domain is a sensor of granular pH that controls the exocytotic machinery. J Cell Biol 203, 283–298.

Ran, F.A., Hsu, P.D., Wright, J., Agarwala, V., Scott, D.A., and Zhang, F. (2013). Genome engineering using the CRISPR-Cas9 system. Nat. Protocols 8, 2281–2308.

Ren, Y., Li, C., Feng, L., Pan, W., Li, L., Wang, Q., Li, J., Li, N., Han, L., Zheng, X., et al. (2016). Proton Channel Activity of Influenza A Virus Matrix Protein 2 Contributes to Autophagy Arrest. J. Virol. 90, 591–598.

Rueden, C.T., Schindelin, J., Hiner, M.C., DeZonia, B.E., Walter, A.E., Arena, E.T., and Eliceiri, K. W. (2017). ImageJ2: ImageJ for the next generation of scientific image data. BMC Bioinformatics 18, 529.

Sanjana, N.E., Shalem, O., and Zhang, F. (2014). Improved vectors and genome-wide libraries for CRISPR screening. Nat Meth 11, 783–784.

Sanjuan, M.A., Dillon, C.P., Tait, S.W.G., Moshiach, S., Dorsey, F., Connell, S., Komatsu, M., Tanaka, K., Cleveland, J.L., Withoff, S., et al. (2007). Toll-like receptor signalling in macrophages links the autophagy pathway to phagocytosis. Nature 450, 1253–1257.

Schindelin, J., Arganda-Carreras, I., Frise, E., Kaynig, V., Longair, M., Pietzsch, T., Preibisch, S., Rueden, C., Saalfeld, S., Schmid, B., et al. (2012). Fiji: an open-source platform for biological-image analysis. Nature Methods 9, 676–682.

Shalem, O., Sanjana, N.E., Hartenian, E., Shi, X., Scott, D.A., Mikkelsen, T.S., Heckl, D., Ebert, B.L., Root, D.E., Doench, J.G., et al. (2014). Genome-Scale CRISPR-Cas9 Knockout Screening in Human Cells. Science 343, 84–87.

Shirakawa, R., Fukai, S., Kawato, M., Higashi, T., Kondo, H., Ikeda, T., Nakayama, E., Okawa, K., Nureki, O., Kimura, T., et al. (2009). Tuberous Sclerosis Tumor Suppressor Complex-like Complexes Act as GTPase-activating Proteins for Ral GTPases. J. Biol. Chem. 284, 21580–21588.

Sumner, J.-P., Dow, J.A.T., Earley, F.G.P., Klein, U., Jäger, D., and Wieczorek, H. (1995). Regulation of Plasma Membrane V-ATPase Activity by Dissociation of Peripheral Subunits. J. Biol. Chem. 270, 5649–5653.

Syrjä, P., Anwar, T., Jokinen, T., Kyöstilä, K., Jäderlund, K.H., Cozzi, F., Rohdin, C., Hahn, K., Wohlsein, P., Baumgärtner, W., et al. (2017). Basal Autophagy Is Altered in Lagotto Romagnolo Dogs with an ATG4D Mutation. Vet Pathol 54, 953–963.

Tchasovnikarova, I.A., Timms, R.T., Matheson, N.J., Wals, K., Antrobus, R., Göttgens, B., Dougan, G., Dawson, M.A., and Lehner, P.J. (2015). Epigenetic silencing by the HUSH complex mediates position-effect variegation in human cells. Science 348, 1481–1485.

Wit, E. de, Spronken, M.I.J., Bestebroer, T.M., Rimmelzwaan, G.F., Osterhaus, A.D.M.E., and Fouchier, R.A.M. (2004). Efficient generation and growth of influenza virus A/PR/8/34 from eight cDNA fragments. Virus Research 103, 155–161.

Xu, Y., Zhou, P., Cheng, S., Lu, Q., Nowak, K., Hopp, A.-K., Li, L., Shi, X., Zhou, Z., Gao, W., et al. (2019). A Bacterial Effector Reveals the V-ATPase-ATG16L1 Axis that Initiates Xenophagy. Cell 178, 552–566.e20.

Yu, Z.-Q., Ni, T., Hong, B., Wang, H.-Y., Jiang, F.-J., Zou, S., Chen, Y., Zheng, X.-L., Klionsky, D.J., Liang, Y, et al. (2012). Dual roles of Atg8-PE deconjugation by Atg4 in autophagy. Autophagy 8, 883–892.

Zhirnov, O.P., and Klenk, H.D. (2013). Influenza A Virus Proteins NS1 and Hemagglutinin Along with M2 Are Involved in Stimulation of Autophagy in Infected Cells. Journal of Virology 87, 13107–13114.

